# Fully flexible molecular alignment enables accurate ligand structure modelling

**DOI:** 10.1101/2023.12.17.572051

**Authors:** Zhihao Wang, Fan Zhou, Zechen Wang, Yong-Qiang Li, Sheng Wang, Liangzhen Zheng, Weifeng Li, Xiangda Peng

## Abstract

Accurate protein-ligand binding poses are the prerequisites of structure-based binding affinity prediction, and also provide the structural basis for in depth lead optimization in small molecule drug design. Ligand-based modeling approaches primarily extract valuable information from the structural features of small molecules to assess their potential as drug candidates against specific targets. However, it is challenging to provide reasonable predictions of binding poses for different molecules, due to the complexity and diversity of the chemical space of small molecules. Similarity-based molecular alignment techniques can effectively narrow the search range, as structurally similar molecules are likely to have similar binding modes, with higher similarity usually correlating to higher success rates. However, molecular similarity isn’t consistently high because molecules often require changes to achieve specific purposes, leading to reduced alignment precision. To address this issue, we propose a new alignment method—Z-align. This method uses topological structural information as a criterion for evaluating similarity, reducing the reliance on molecular fingerprint similarity. Our method has achieved significantly higher success rates than other methods at moderate levels of similarity. Additionally, our approach can comprehensively and flexibly optimize bond lengths and angles of molecules, maintaining high accuracy even when dealing with larger molecules. Consequently, our proposed solution helps in achieving more accurate binding poses in protein-ligand docking problems, facilitating the development of small molecule drugs.

## 1 Introduction

The key challenge in small molecule drug design lies in determining precise protein-ligand binding patterns and binding strength [1, 2]. The accuracy of the predicted structure models would significantly impact the molecule optimization directions in lead optimizations [3]. Different types of computational methods have been proposed to predict protein-ligand binding structures, and numerous researchers have also developed various scoring functions to evaluate the strength of these interactions [4, 5].

Molecular docking is one of the most commonly used methods to predict the protein-ligand complex structures [6]. This method samples a wide range of conformations of ligand and/or side chain on protein and then evaluates the quality of the binding mode through a set of scoring functions [7, 8].

Traditional scoring functions estimate the binding affinity or calculate the binding free energies of the complexes using either knowledge-based parameterization, statistical modeling [7, 8, 9, 10], or force field-based functions [11]. In addition, a lot of the machine learning-based or deep learning-based scoring functions [12, 13, 14, 15, 16, 17, 18, 19] have been developed in recent years. These models are generally used as a post-scoring method to re-score the poses of the ligand generated by various docking applications.

When performing molecular docking, a large number of ligand poses (and sometimes side-chain orientations of proteins in flexible docking protocols [20]) are sampled guided by the scoring functions. Representative sampling algorithms include the Monte Carlo method in AutoDock Vina [21] or genetic algorithm in GOLD [22]). They all require a large computational time to find a correct global minimum. Given that the scoring functions may not perfectly describe the protein-ligand interactions on the entire potential energy surface, the expectations of a global minimum of the protein-ligand binding pattern thus could not be guaranteed, leading to many false positive predictions [23].

Interestingly, similar situations are encountered in early protein structure prediction problems. Before the emergence of AlphaFold2 [24], various methods were attempted to predict protein structures, including residue contact map-based structure predictions [25, 26], ab initio molecular dynamics simulations [27], and others. However, the accuracy of these models was quite limited, often due to difficulties in sampling and the insufficiency of force field precision. Another clever traditional approach to solving protein structure problems was based on the use of templates. With the assumption that similar proteins obey similar structures, researchers have been able to reproduce the protein structures more precisely by relying on protein structure templates from the PDB database [28, 29].

Currently, when predicting the structures of protein-ligand complexes, researchers are employing similar strategies, template-based methods, with success in the ligand prediction category of the Critical Assessment of Structure Prediction 15 (CASP15) competition [30, 31, 32]. This method consists of two key steps. The first step is to find the template, which has a homologous sequence and a crystal structure with the co-crystal substrate. The second step is to model the protein-ligand conformation based on the co-crystal substrate (template molecule). This process is also essentially a flexible structural alignment. Compared to molecular docking, template-based modeling based on prior knowledge may reduce false positives and greatly narrow down the conformational space.

Structure alignment involves comparing molecular structures to identify segments that are similar enough to be aligned [33]. By aligning molecules to existing co-crystal structures, the newly obtained protein-ligand binding conformations are likely to be native or near-native. Presently, methods for aligning small molecules fall into two main categories: rigid alignment and flexible alignment [34]. Rigid alignment methods primarily rely on shape-based techniques [35, 36, 37]. These methods often involve fitting the Gaussian function to the molecular volume to maximize the overlap of molecular shape. However, there are critical limitations that the query ligands cannot transform their geometric structures to match the template molecules, nor can they accommodate different shapes of binding pockets. In contrast, flexible alignment largely overcomes these limitations. It starts with the identification of structurally or chemically equivalent features on the atomic level and then proceeds by rotating rotatable bonds enabling geometric transformation that significantly enhances the alignment precision [34, 38, 39].

Nonetheless, alignment strategies that solely involve rotating bonds without adjusting bond lengths do not offer a genuine “fully flexible” alignment. This limitation might not have distinct impacts on smaller molecules, but for larger molecules, especially those with elongated structures, errors in bond lengths can magnify, leading to a significant reduction in alignment precision. Moreover, in practical applications, the degree of similarity between the template and query molecules is inherently limited, complicating the identification of equivalent atomic pairs. Consequently, when the similarity is moderate, which is the most common in practical scenes, the existing methods often fail to deliver satisfactory outcomes.

In this study, we propose a novel fully flexible small molecule alignment method, named Z-align. Z-align avoids the strict equivalent atomic pair searching procedure. When the similarity between molecules is not high, the rational construction of equivalent atomic mapping is challenging. Instead, Z-align directly places the query molecule in a potential field constructed from a template molecule, where spatial overlap or chemical similarity will be rewarded. After sampling the overall conformation and rotatable bonds, multiple conformations of the query molecule undergo energy minimization with the potential field mentioned above and the constraints of the molecule’s all-atom force field (to ensure reasonable geometric properties). In this way, Z-align can adjust both bond lengths and bond angles while ensuring the structural rationality of the molecule. In testing, our method demonstrates notable advantages. In self-alignment tasks, it can align almost all ligands to their native poses successfully and exhibit the best performance in scenarios with medium/low similarity levels.

## 2 Methods

### 2.1 Overview of Z-align

The workflow of Z-align is illustrated in Figure 1**A**. The whole process can be divided into three steps: force field construction, interaction potential application, and sampling with energy minimization. In step 1, the algorithm loads the input files and generates corresponding force field information. In step 2, the attractive and repulsive potential will be applied to similar query-template and grid-query atomic pairs which are determined by the force field information. Then, the initial query ligand is translated and rotated in three-dimensional space for sampling. Meanwhile, the rotational sampling of chemical groups connected by rotatable bonds is also conducted. At last, after energy minimization performed by L-BFGS algorithm provided by OpenMM [40], the pose with the lowest energy will be taken as the output conformation.

**Figure 1:**
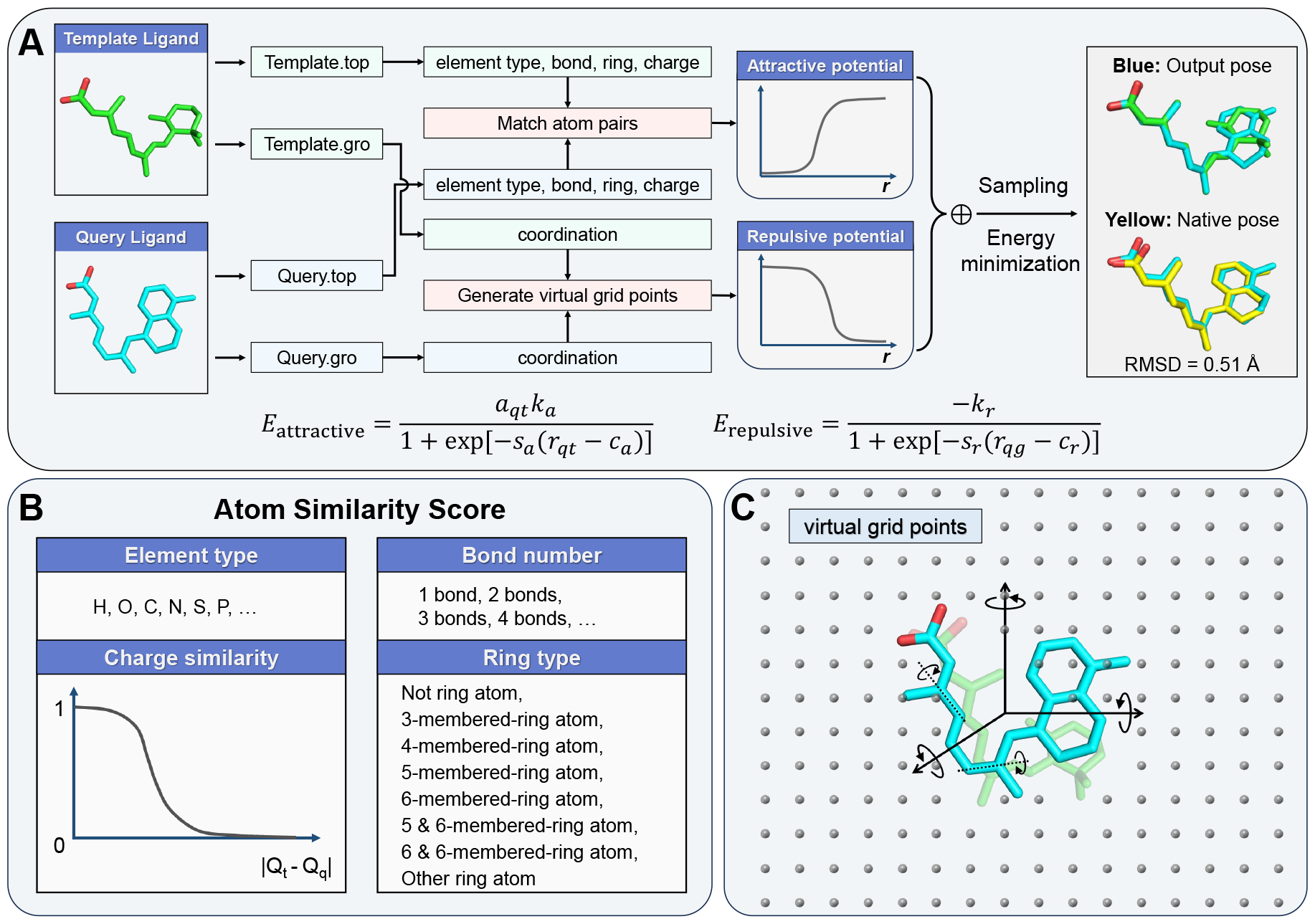
(**A**)The workflow of Z-align. (**B**)Atomic indicators for finding similar atomic pairs. (**C**)Virtual grid points construction diagram.

### 2.2 Force field construction

The 3D structures of the query ligand and the template ligand are first processed by OpenBabel [41] in order to add the missing hydrogen atoms. The processed structures are then used as the input files for ACPYPE [42] to generate suitable force field with GAFF2 [43]. The bond lengths inside the molecule are constrained by the force field using the spring potentials, which keeps the molecule in a reasonable structure. Flexible constraints on bond lengths and bond angles make it possible for small molecules to undergo a certain degree of deformation, which is beneficial for the alignment as it brings about more sampling of conformations.

### 2.3 Interaction potential

#### 2.3.1 Attractive potential

The most import step in molecule alignment should be the atom pair matching between template ligand and query ligand. Different from LS-align [34] and FitDock [38], we use the chemical and topological information to find similar atomic pairs instead of detailed hierarchical strategy to find equivalent atomic pairs. The interaction parameters in the force field can provide basic information about the geometrical and elemental properties of small molecules. Here, the element types, charge values, bond numbers and ring types were chosen as similarity indicators. The similarity score is defined as

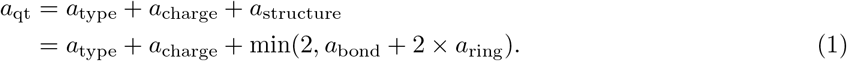

As shown in Figure 2, the values in matrix are set to one or zero if the elements are identical or not. The topological structure information of an atom is evaluated by the number of chemical bonds it forms and whether it is an atom in a ring. If the query atom *i* and the template atom *j* form the same number of bonds, the corresponding matrix element will be set to 1. At the same time, the algorithm will automatically determine the ring information, which includes eight types, as illustrated in Figure 1**C**. We considered that the ring type carries more important topological information than the bonding numbers, so its weight is twice that of the bonding similarity. Moreover, in order to maintain a balance between topological and chemical information, the score for structural similarity is capped at a maximum of two points. The charge similarity score is determined by a sigmoid-like function whose shape is plotted in Figure 1**B**. The closer the values of the charges are, the closer the similarity score is to 1:

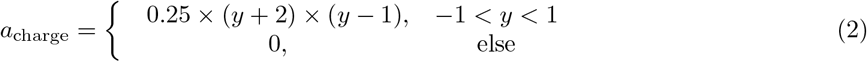

where *y* = 10 × (|*Q*_t_ − *Q*_q_| − 1). *Q*_t_ and *Q*_q_ are the charges of the atoms in template and query molecules.

**Figure 2:**
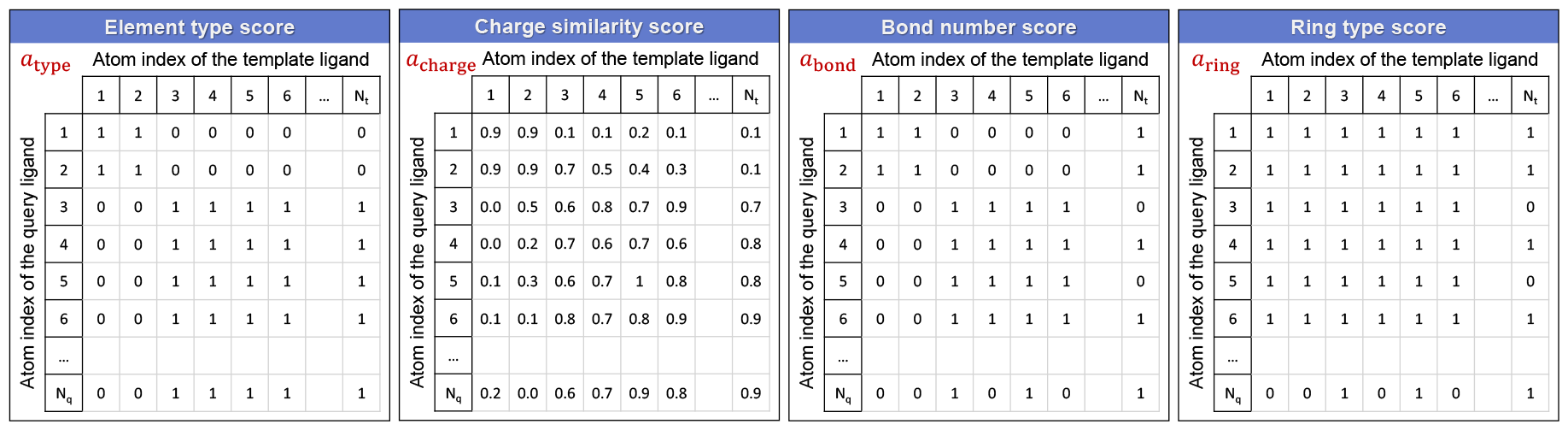
Atom similarity matrix of element type, bond number, ring type and charge values.

The attractive potential will be applied to the query-template atomic pairs *ij* if their summed similarity score 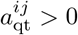. The potential function is defined as

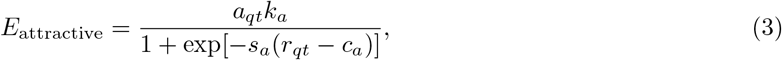

where

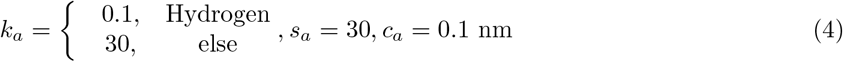

are several selected constants. The *r*_*qt*_ in equation (2) is the distance between two similar atoms after removing the coordinate difference of geometric centers of the query and template molecules. For different atomic pairs (for example, pair *ij* and pair *ik*), the similarity score 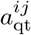 and 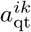 are different, which makes the strength of the attractive potential different. Hence, a greater resemblance between atoms results in a heightened attraction force between them.

The reason why we use the sigmoid-like formula as the attractive potential is that we need it to provide switching between presence and absence, and that it can flexibly control the force range. We intentionally keep the attractive interaction range very short to avoid complex conflicts with other functions. Therefore, the attractive potential only works when pairs of atoms are very close (*<* 0.5 Å), bringing them closer even further.

#### 2.3.2 Repulsive potential

In addition to the attractive potential between pairs of similar query and template atoms, we also apply a repulsive potential to all atoms in the query ligand. As shown in Figure 1**C**, Z-align will construct a uniform distribution of virtual grid points in the surrounding space with each atom of template ligand (the green molecule) as the center. The difference of geometric centers of the query and template ligands are removed by translating the query ligand. The conformation of the query molecule after translation may clash with the virtual grid points around the template molecule. The repulsive potentials applied between the grid points and query atoms will “push” the molecule away from the grid to achieve maximum overlap between template and query ligands. The repulsive potential shares a similar form with the attractive potential, which is written as

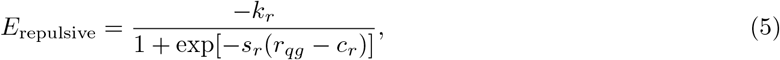

where

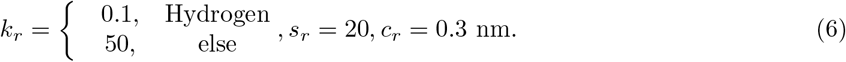

The *r*_*qg*_ in equation (5) is the distance between query atoms and grid points.

Different from the attractive potential, the sigmoid function for the repulsive potential has a longer interaction range, which helps to construct a smooth potential energy surface to reduce local traps.

### 2.4 Sampling and energy minimization

The global sampling is performed by rotations along the three principal axes, while rotational sampling of the rotatable bonds is also conducted simultaneously. Each conformation generated during sampling will serve as an initial structure input into OpenMM [40] for optimization using molecular dynamics simulations to minimize the total energy. The total energy consists of three parts:

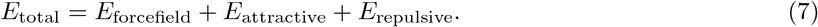

The force field part is set to be dimensionless, which can be expanded as

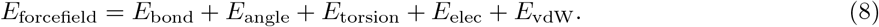

Finally, the conformation with the lowest energy will be outputted.

### 2.5 Datasets

The benchmark data sets were divided into two parts for the purpose of “self alignment” and “cross-docking alignment”.

For self alignment, the native structures were taken as the template ligands and the query ligands were generated randomly by RDKit [44] with the native poses as the input files. We calculated the root mean square deviations (RMSD) of the aligned poses relative to the native pose for evaluation using spyrmsd [45]. The datasets employed here are the refined set of PDBbind v2018 [46] and the cross-docking database 3D Disco [47]. As for the 3D Disco dataset, the receptor structures were acquired from DUD-E dataset [48]. Take the receptor structure that shares the same PDB entry with the ligand (representative ligand) as representative receptor. Then superimpose other ligands from the same target to the representative receptor. The conformations here were assumed to be correct or near-native thus can be used as “native” poses. In self alignment, we put “native” poses into RDKit to obtain random poses. And then we aligned the random poses to the “native” poses that serve as the templates. The initial RMSD distributions of the datasets were plotted in Figure S1.

In terms of cross-docking alignment, 3D Disco was also selected as the benchmark. Different from self alignment, the template molecules were the representative ligands and the query molecules were the random conformations generated from “native” poses of same targets. The success rate of molecular alignment usually increases with the similarity between query molecules and template molecules. Thus, we assessed the Morgan fingerprint similarity by RDKit between query molecules and template molecules. As shown in Figure S2, the query-template molecular pairs are mainly distributed in the regions with low similarity.

Besides, we made up a dataset called large molecule for additional evaluation of self alignment. This dataset consists of 10415 molecules with each molecule having more than 30 but no more than 100 heavy atoms. Due to the excessive number of atoms and rotatable bonds in this dataset, many randomly generated structures by RDKit are not reasonable. Therefore, we use the OpenBabel software package for the randomization of initial structures.

Due to variations in the number of molecules that different methods can successfully process and output, for a fair comparison, we chose the common molecules that all algorithms can successfully output when calculating success rates. The number of molecules successfully output by each alignment method is summarized in Tables S1 and S2.

### 2.6 Other methods used for comparison

Three alignment programs were chosen to make comparison.

LIGSIFT is an open-source alignment method that utilizes Gaussian functions to model the shape of the entire molecule instead of analyzing at the atomic level [37]. Metropolis Monte Carlo simulation is used to realize the structure overlap of template molecules and query molecules. Please note that the alignment process in LIGSIFT is rigid without any flexible movements.

LS-align employs both rigid and flexible alignment techniques, incorporating inter-atom distance, mass, and chemical bond comparisons [34]. It can achieve fast and accurate structural alignment through iterative heuristic search. The flexible mode Flexi-LS-align was used here for testing.

FitDock can serve for both molecular alignment and molecular docking purposes [38]. By hierarchical multi-feature comparison, it can achieve accurate matching of equivalent atomic pairs and substructures and finally outputs refined docking poses.

## 3 Results and discussion

### 3.1 Z-align and other alignment methods with flexible modules performed well in self alignment task

We performed self-align tests on the refined set of PDBbind v2018 and 3D Disco. The success rates of each method in different RMSD cutoff ranges are shown in Figure 3. Z-align could align almost all query ligands within RMSD of 2 Å, showing marvelous performance. The success rates of aligment within 2 Å were 99.73% and 98.63%, respectively. In self-align tasks where the chemical formulas of the template and query molecules are identical, the element type, bond count, and ring configuration of an atom in the query molecule match those of its counterpart in the template molecule. Thus our method demonstrates excellent performance. Similarly, when the chemical formula is exactly the same, the FitDock method can accurately find the equivalent atomic pair using hierarchical multi-feature atomic pair information comparison, achieving success rates of 99.68% and 99.81% within a 2 Å range, demonstrating the best performance. Among the four alignment methods assessed, the three with flexible alignment exhibited notably superior performance compared to LIGSIFT, which was limited to rigid alignment only.

**Figure 3:**
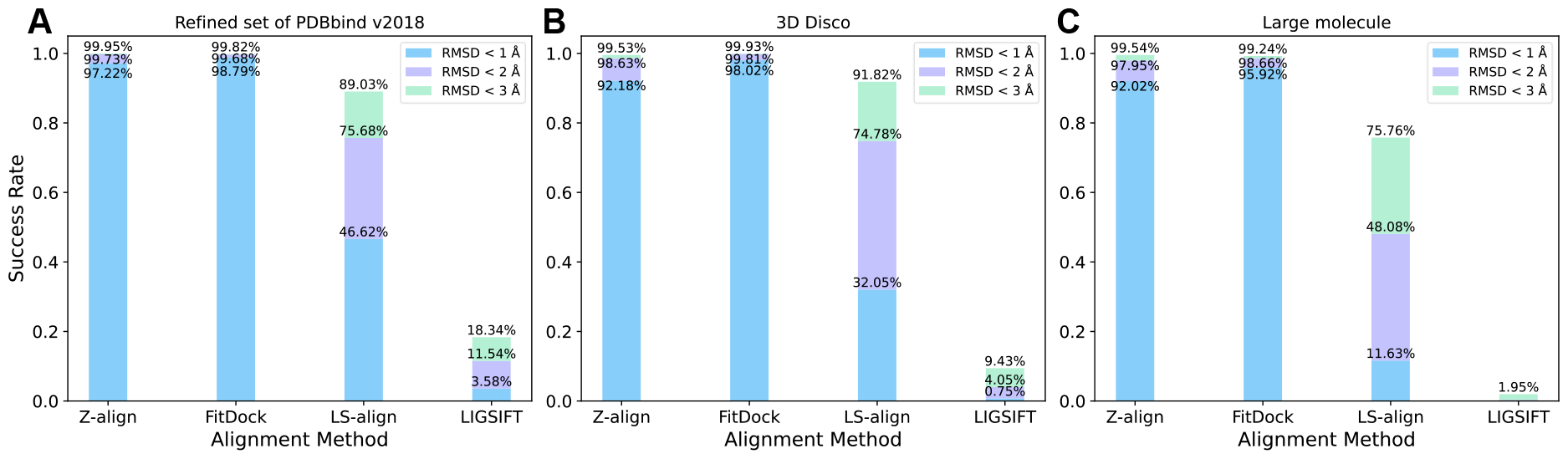
The self alignment success rates of Z-align, FitDock, LS-align and LIGSIFT at the refined set of PDBbind v2018 (**A**), 3D Disco dataset (**B**) and Large molecule (**C**).

### 3.2 Z-align outperformed in cross-docking alignment task

The results of cross-docking can be used as an important indicator to assess an alignment method. Because different ligands binding to the same pocket are likely to follow similar binding patterns, and cross-docking serves as an important approach to model unknown binding patterns based on known binding conformations. This has guiding significance for the design of novel drugs. If we have a ligand known to bind to this pocket and obtain a sufficiently accurate co-crystal structure, we can then use this ligand as a reference to adjust the structure of the new molecule. Hence, the value of an alignment method lies in its ability to exhibit good performance in cross-docking tasks.

As shown in Figure 4**A**, the outcomes of cross-docking alignment significantly differ from those of self-alignment, showcasing notable decreases in success rates compared to the self-alignment results. Z-align manifests the best performance among the tested four methods. However, the success rate (RMSD *<* 2Å) of FitDock is only 13.52% which is not even as good as that of LS-align (15.06%). The main reason of this difference can be attributed to the discrepancy between the query molecules and the template molecules. Due to the diminished similarity, the stringent atom pair matching approach does not perform as effectively as it does in the self-alignment task.

**Figure 4:**
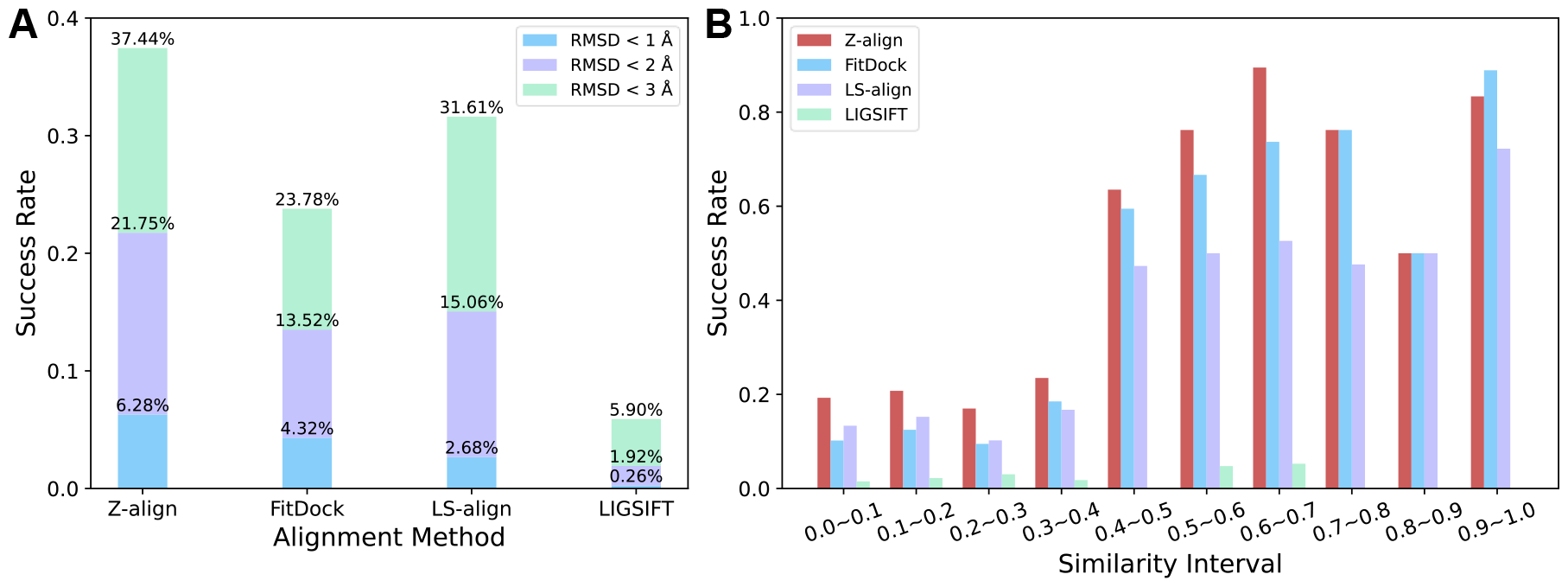
(**A**) The cross-docking alignment success rates of Z-align, FitDock, LS-align and LIGSIFT at the 3D Disco dataset. (**B**) The cross-docking alignment success rates of Z-align using templates within different similarity intervals.

For verification, we further calculated the success rates at different similarity intervals, which was plotted in Figure 4**B**. FitDock has the highest success rates when the similarity exceeds 0.9, yet it experiences the most notable decrease in success rates as the similarity diminishes. In practice, molecules of moderate similarity are of greater importance. Because it is expected that the drug molecule has a certain similarity with the template molecule to ensure a similar binding strength, it also needs some structural differences in the drug molecule to achieve a certain specific biological function. Z-align maintains consistently high success rates within high similarity intervals and excels particularly well when the similarity is between 0.4 and 0.8. This is because although the query and template ligand are different, there are always some analogous topological structures (for example, similar ring configurations) in the molecules. Our method can detect this structural similarity and then apply appropriate attractive potentials to the similar atomic pairs. Moreover, the repulsive potentials introduced by grid points will restrain the whole molecule into a confined space which prevents arbitrary expansion of the molecule. Additionally, the alignment process is conducted by molecular dynamics, where the force field parameters also impose proper constraints on the molecule configurations avoiding unreasonable deformation. Please note that at the interval of 0.8∼0.9, only 2 query-template molecule pairs were found in the dataset (Figure S2). Therefore, Z-align, FitDock, and LS-align achieved the same success rate of 0.5 as they all aligned only one ligand with 2 Å.

### 3.3 Z-align improved the optimization of bond length and bond angle of large molecules

Molecules with a greater number of atoms can assess the ability of alignment methods to handle complex problems. The complexity of molecules in the Large molecule dataset has notably increased the difficulty of alignment, resulting in a significant decrease in the proportion of molecules that each algorithm can successfully process (as shown in Table S1). We selected 7594 molecules that can be processed by all algorithms for comparison of success rates (Figure 3**C**). Z-align and FitDock also manifest high success rates in such hard dataset while LS-align and LIGSIFT show apparent declines.

We further analyzed the bond lengths and bond angles of the molecules output by different algorithms and compare them with the template molecules which were shown in Figure 5. Here, only chemical bonds without hydrogen atoms were taken into consideration. As LIGSIFT only deploys rigid alignment which means the lengths and angles didn’t change during processing, we didn’t do such analyses on it. The results show that Z-align can improve the correlation coefficients of both bond length and bond angle. The force field based alignment imposes proper restraint on the bond lengths according to the bond types. For some similar chemical bonds, our method applied similar harmonic potentials to them, limiting the bond lengths to fluctuate around several certain values. Therefore, several distinct clusters can be found in Figure 5**B**, exhibiting a staircase-like pattern. Different from the other methods, the distribution of the bond length is not continuous, but exhibits several gaps. Thus, Z-align is able to identify irregular bond lengths and subsequently adjust them to fall within appropriate ranges. There are no distinct improvements in bond length correlations of FitDock and LS-align as their frameworks lack the capability to modify bond lengths. Similar to the bond length distribution, there are also several clusters in bond angle distribution on account of the restraint of the force field. The restraint on bond angles brought an big improvement of correlation coefficient (0.136, 18.48%). FitDock exhibits clear linear correlation between template bond angles and output bond angles, which gave the value of 0.949. As for LS-align, the correlations of bond length and bond Angle are not improved, resulting in a low success rate of only 41% within 2 Å.

**Figure 5:**
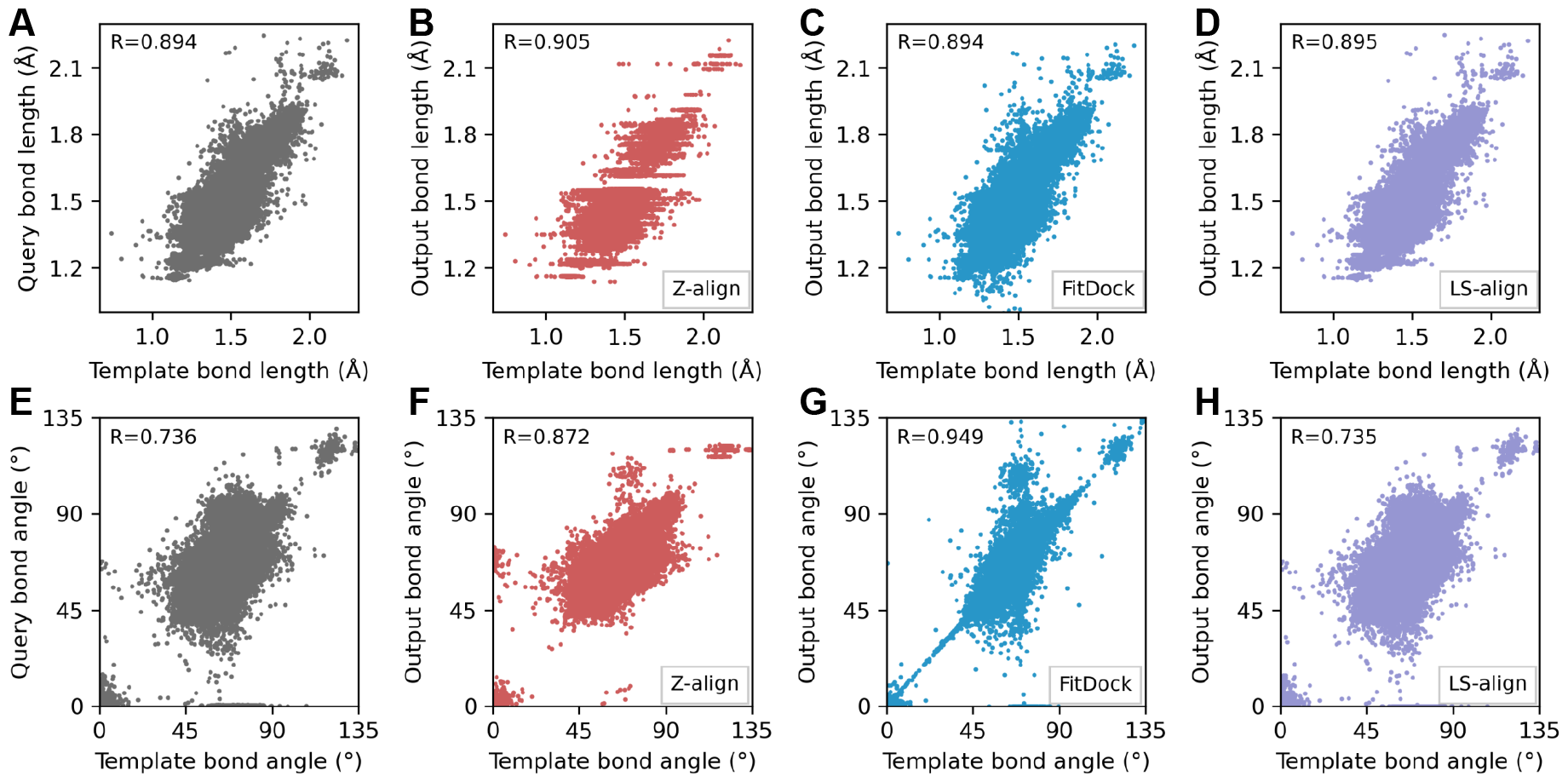
Comparison between the bond length (**A**) and bond angle (**E**) of the template and query molecules in large molecule dataset. Comparison between the bond length (**B-D**) and bond angle (**F-H**) of the template and output molecules aligned by Z-align (**B, F**), FitDock (**C, G**) and LS-align (**D, H**) in large molecule dataset. The Pearson correlation coefficients (R) were also illustrated in each sub figure.

### 3.4 Case study

As mentioned above, adjusting bond lengths and bond angles contributes to improving alignment accuracy. To illustrate this directly, three elongated chain-like molecules, adenosine triphosphate (ATP), nicotinamide adenine dinucleotide (NAD), and Coenzyme A (COA), were selected as evaluation cases shown in Figure 6**A**-**C**. The common characteristic among these three molecules is that they all contain an adenosine moiety and a ribose, and feature elongated chains composed of phosphate groups. The compared three methods was able to identify the adenosine moieties of molecules and aligned them in nearly correct manners. However, starting from the ribose, conformations obtained from LS-align deviate from the template molecule. These deviations progressively accumulate as the molecules elongate, eventually resulting in a substantial difference of the overall conformations from the template molecules. We also took some output poses from cross-docking task as instances (Figure 6**D**-**F**). The reliance on structural similarity varies distinctly among the three algorithms. For LS-align and FitDock, the query ligands were superposed to the template ligands in a reasonable manner. Nevertheless, at low similarity levels, certain functional groups were oriented incorrectly on account of mismatches in equivalent substructures. Consequently, the employment of highly flexible alignment framework significantly enhances the precision of molecular modeling.

**Figure 6:**
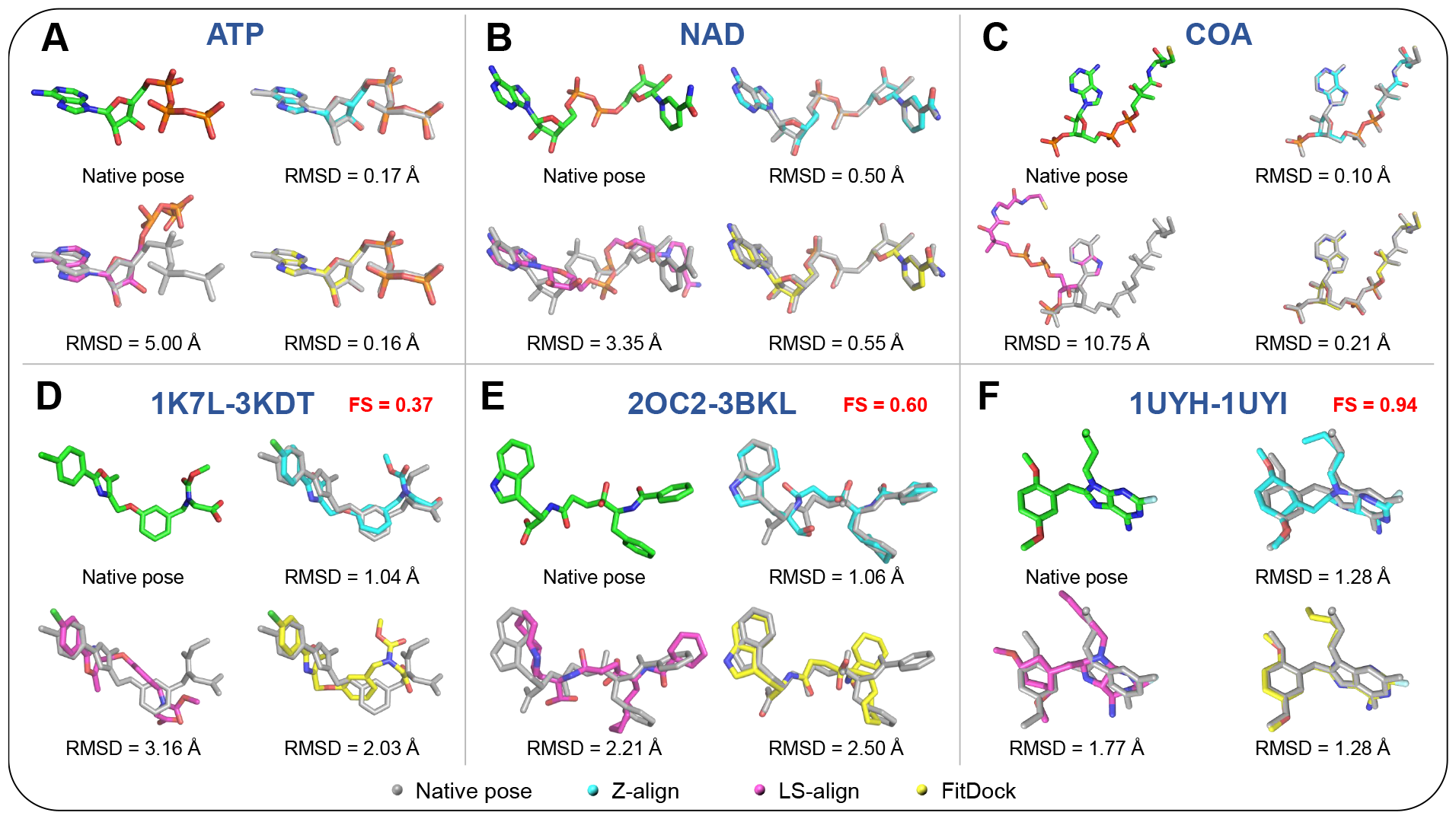
Examples of molecules output by Z-align, FitDock and LS-align. The large molecules selected here are ATP (**A**), NAD (**B**), and COA (**C**), respectively. (**D**)-(**F**) are the molecules selected from the cross-docking test. 1K7L-3KDT (2CO2-3BKL, 1UYH-1UYI) stands for the PDB code of the template-query ligand pairs. The fingerprint similarity (FS) between template-query ligand pairs were shown in each panel.

### 3.5 Alignment speed evaluation

We randomly selected 1000 ligands from the refined set of PDBbind v2018 to perform the alignment speed test. The methods that employ flexible alignment function were tested for comparison. The average time for an alignment task of Z-align, FitDock and LS-align are given by 698.75 s, 6.01 s and 0.55 s, respectively. As stated, Z-align takes the most time for a single alignment task. It can be attributable to the time-consuming charge calculation in the step of force field construction. Although Z-align takes a lot of time to construct the force field, it will be convenient if molecular dynamic simulations are needed later. We also analyzed the relationship between the alignment time and the number of rotatable bonds, as shown in Figure S3. Owing to some very time-consuming tasks, the time of FitDock is not as short as reported [38]. And for these complicated tasks (for example, the PDB IDs of 1fkb, 4l6t, 4gah and 3old), it generated a significantly larger number of conformations, resulting in a dramatic increase in memory usage that eventually exceeded the capacity and terminated without giving any output pose. Therefore, overall, each alignment framework has its advantages and shortcomings. We still need to invest more effort to further enhance their comprehensive capabilities.

## 4 Conclusion

Molecular alignment, driven by structural similarities, holds significant value in solving protein-ligand docking issues by narrowing down the ligand conformational space. This approach aids in predicting more accurate binding poses, thereby reducing false positives during virtual screening. In this work, we developed a molecular alignment method for optimizing ligand poses based on template structures. Importantly, our method bypasses equivalent atomic pair matching, focusing instead on the chemical and structural similarities of atomic pairs. This strategy offers distinct advantages, particularly in scenarios with moderate or even low molecular similarity in cross-docking alignment tasks, where Z-align manifests superior performance. Additionally, our method incorporates bond length and bond angle optimization, establishing a fully flexible alignment framework that effectively addresses cases involving large molecules. Hence, in practical virtual screening scenarios, this fully flexible alignment method, which relies less on molecular similarity, is anticipated to demonstrate greater advantages.

## 5 Acknowledgement

This work is financially supported by the Natural Science Foundation of Shandong Province (ZR2020JQ04) and Local Science and Technology Development Fund Guided by the Central Government of Shandong Province (YDZX2022089).

