## Supporting information Figure S1 for "Fully flexible molecular alignment enables accurate ligand structure modelling"

Part 1. The initial RMSD distributions of the datasets for self-alignment task and the similarity distribution of 3D Disco.

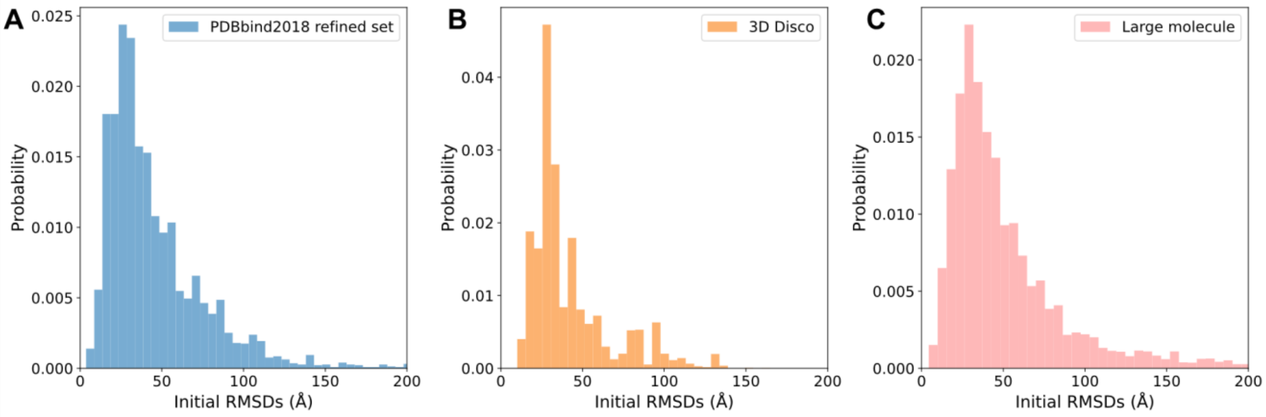


Figure S1. The initial RMSD distributions of PDBbind 2018 refined set, 3D Disco and large molecule.

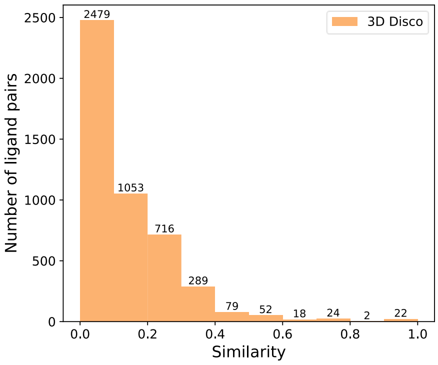


Figure S2. The similarity distributions of 3D Disco.

Part 2. The numbers of molecules of different datasets.

Table S1. The numbers of molecules of different datasets in self-alignment task. The numbers of molecules randomly generated by RDKit or OpenBabel are indicated in parentheses after the names of the datasets. “Output” means the number of molecules that can be successfully output by a certain alignment method. “Common” refers to the number of common molecules that can be output by all methods.

| | **Refined set of PDBbind v2018 (4463)** | | | | **3D Disco Self-alignment (4826)** | | | | **Large molecule (10415)** | | | |
|---|---|---|---|---|---|---|---|---|---|---|---|---|
| Num of molecules | Z-align | FitDock | LS-align | LIGSIFT | Z-align | FitDock | LS-align | LIGSIFT | Z-align | FitDock | LS-align | LIGSIFT |
| Output | 4411 | 4437 | 4463 | 4457 | 4261 | 4801 | 4826 | 4824 | 7759 | 8574 | 8805 | 9266 |
| Common | 4384 | 4384 | 4384 | 4384 | 4243 | 4243 | 4243 | 4243 | 7594 | 7594 | 7594 | 7594 |
| RMSD < 1 Å | 4262 | 4331 | 2044 | 157 | 3911 | 4159 | 1360 | 32 | 6991 | 7286 | 883 | 1 |
| RMSD < 2 Å | 4372 | 4370 | 3318 | 506 | 4185 | 4235 | 3173 | 172 | 7441 | 7495 | 3651 | 25 |
| RMSD < 3 Å | 4382 | 4376 | 3903 | 804 | 4223 | 4240 | 3896 | 400 | 7562 | 7539 | 5753 | 148 |

Table S2. The numbers of molecules of 3D Disco dataset in cross-docking alignment task.

| | **3D Disco Cross-docking alignment (4735)** | | | |
|---|---|---|---|---|
| Num of molecules | Z-align | FitDock | LS-align | LIGSIFT |
| Output | 4266 | 4713 | 4734 | 4698 |
| Common | 4217 | 4217 | 4217 | 4217 |
| RMSD < 1 Å | 265 | 182 | 113 | 11 |
| RMSD < 2 Å | 917 | 570 | 635 | 81 |
| RMSD < 3 Å | 1579 | 1003 | 1333 | 249 |

Part 3. Results of alignment speed test.

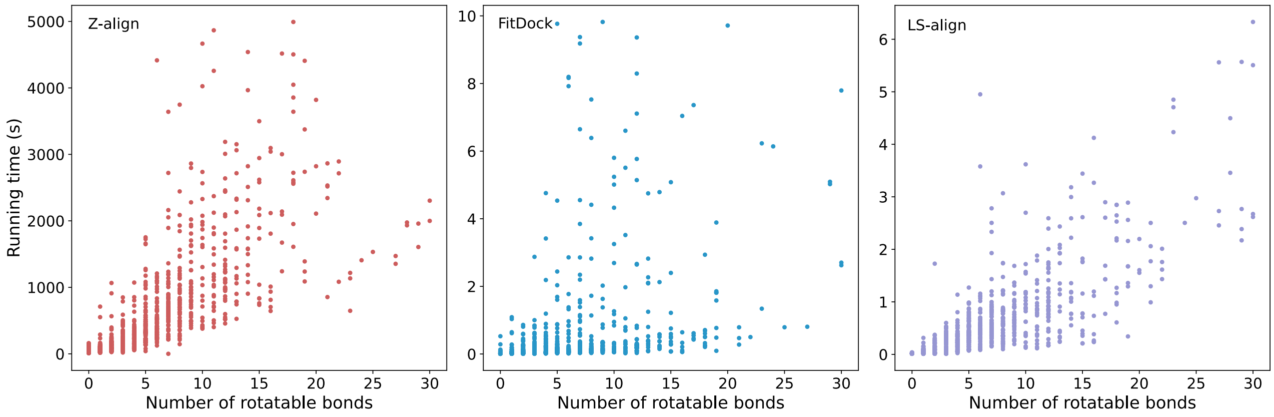


Figure S3. The docking time of Z-align, FitDock and LS-align with respect to number of rotatable bonds.
